# SGIP1 binding to the α-helical *H9* domain of cannabinoid receptor 1 promotes axonal surface expression

**DOI:** 10.1101/2023.07.18.549510

**Authors:** Alexandra Fletcher-Jones, Ellen Spackman, Tim J. Craig, Yasuko Nakamura, Kevin A. Wilkinson, Jeremy M. Henley

## Abstract

Endocannabinoid signalling mediated by cannabinoid type 1 receptors (CB1Rs) is critical for homeostatic neuromodulation of both excitatory and inhibitory synapses. This requires highly polarised axonal surface expression of CB1R, but how this is achieved remains unclear. We previously reported that the *H9* domain in the intracellular C-terminus of CB1R contributes to polarised surface expression by an unknown mechanism. Here we show the *H9* domain binds to the endocytic adaptor protein SGIP1 to promote CB1R expression in the axonal membrane. Overexpression of SGIP1 increases CB1R axonal surface localisation but has no effect on CB1R lacking the *H9* domain (CB1R^ΔH9^). Conversely, SGIP1 knockdown reduces axonal surface expression of CB1R but does not affect CB1R^ΔH9^. Furthermore, SGIP1 knockdown diminishes CB1R-mediated inhibition of presynaptic Ca^2+^ influx in response to neuronal activity. Together, these data advance mechanistic understanding of endocannabinoid signalling by demonstrating that SGIP1 interaction with *H9* underpins axonal CB1R surface expression to regulate presynaptic responsiveness.

## Introduction

The endocannabinoid system (ECS) is a negative feedback system that homeostatically controls neurotransmission in the brain. By mediating activity-dependent suppression of presynaptic release, the ECS modulates synaptic strength and plasticity, which are fundamental for many brain processes including cognition, appetite/energy expenditure, and learning and memory (1). Moreover, the ECS plays key roles in attenuating stress-induced glutamate release and is implicated in a wide range of neurological and neurodegenerative diseases (2, 3).

Because ECS pharmacology is complex and pleotropic, drugs that act directly on the system often result in unwanted neurological and psychoactive side effects (4). Given these limitations, increased understanding of the biochemistry and cell biology of the ECS could provide new avenues for therapeutic intervention.

In neurons, the main ECS receptor, cannabinoid type 1 receptor (CB1R), is located predominantly at the axonal membrane (5–14), particularly at the presynaptic terminal (15–17). CB1R activation by endocannabinoids released from the postsynaptic membrane suppresses presynaptic neurotransmitter release via G protein-mediated inhibition of presynaptic voltage-gated Ca^2+^ channels (18) and/or adenylyl cyclase activity (19). Thus, the selective targeting of CB1R to the axonal membrane is crucial to its role in regulating activity at the presynapse, yet how this is orchestrated at a molecular level is poorly defined (20).

We have reported previously that CB1R is preferentially and directly axonally targeted through the secretory pathway and that polarity is maintained, at least in part, by the more rapid endocytosis of CB1Rs from the somatodendritic than the axonal membrane (13). Furthermore, we showed that the 21-residue putative helical *H9* domain in the intracellular C-terminal domain of CB1R (ctCB1R) contributes to the delivery and stabilisation of axonal CB1R (13). However, despite this progress, exactly how *H9* promotes the axonal surface distribution of CB1R remains to be determined.

SH3-containing GRB2-like protein 3-interacting protein 1 (SGIP1) is abundant in brain (21) and preferentially localised to axons and presynaptic terminals (22–24). SGIP1 is an endocytic adaptor protein linked to clathrin-mediated endocytosis (25–30) but its precise role(s) are elusive and may be isoform-dependent since the longer, less abundant isoform, SGIP1α, is capable of membrane tubulation whereas SGIP1 itself is not (31).

SGIP1 has been reported to bind ctCB1R in a yeast 2-hybrid study but the site of interaction on CB1R was not determined (23). Nonetheless, expression studies in HEK293 cells have suggested that SGIP1 interferes with agonist-induced internalisation of CB1R and modulates β-arrestin and GRK3 association and downstream ERK1/2 signalling (23, 32). Moreover, SGIP1 knockout mice display disrupted ECS-dependent behaviours and altered responses to Δ^9^-tetrahydrocannabinol (THC), including reduced anxiety, reduced acute nociception, and increased sensitivity to cannabinoid-induced analgesia, while working memory and exploration remain unaltered (33).

Here we report that SGIP1 binds to the CB1R α-helical *H9* domain and acts to stabilise CB1R at the presynaptic membrane. We show that overexpression of SGIP1 increases CB1R at the axonal plasma membrane, whereas SGIP1 knockdown phenocopies the decreased surface expression observed upon *H9* deletion (CB1R^ΔH9^) and impairs CB1R-mediated modulation of synaptic transmission. These data provide mechanistic understanding of how CB1R polarity is established and maintained by identifying SGIP1 as an important mediator of CB1 axonal surface expression. Moreover, these findings open the possibility that manipulating this interaction could be used to regulate the availability of presynaptic CB1R for potential therapeutic benefit.

## Results

### Cloning of SGIP1β from rat cortical neuronal cultures

Deletion of *H9* reduces CB1R surface expression and increases CB1R endocytosis in primary neurons (13), whereas co-expression of SGIP1 enhances CB1R surface expression in HEK293 cells (23). Based on these observations we wondered if SGIP1 interacts with *H9* to regulate CB1R surface expression. To investigate this possibility, we amplified rat SGIP1 from cDNA derived from mRNA extracted from DIV21 primary cortical neurons and subcloned it into a modified pcDNA3.1 to incorporate an N-terminal FLAG tag.

The isolated sequence corresponded to predicted SGIP1 transcript variant X19 (NCBI Reference Sequence: XM_017593774.2; **Figure 1A**). This 660-amino acid variant differs from the full-length canonical Uniprot entry (i.e., transcript variant X9 NCBI Reference Sequence: XM_017593764.2) by two deletions: a single residue deletion in the membrane phospholipid binding domain (MP; Q34) and a 165-residue deletion in the proline-rich domain (PRD).

**Figure 1:**
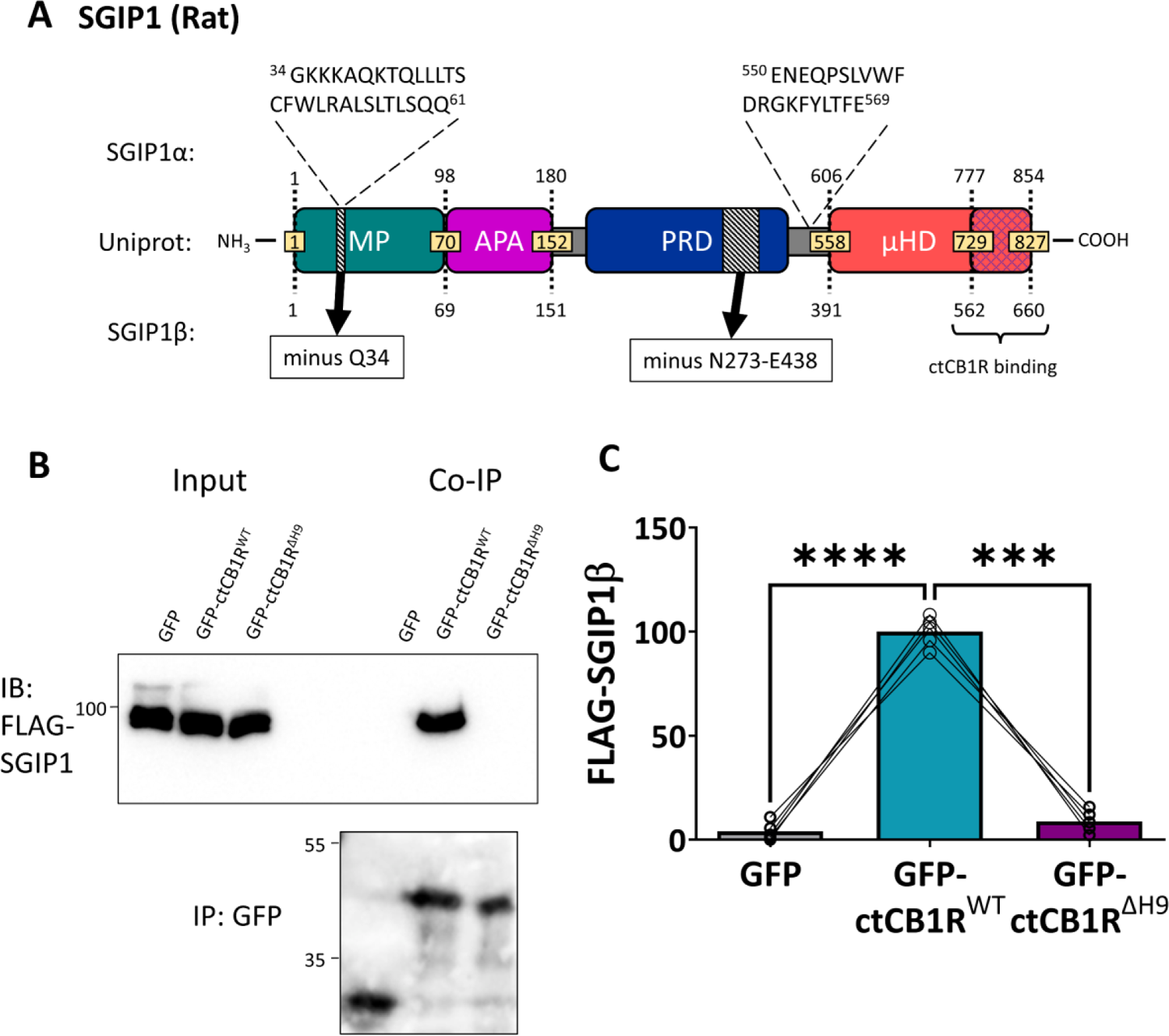
*H9* interacts with SGIP1. (A) Schematic comparing the three known SGIP1 isoforms. The SGIP1 isoform we cloned from a cDNA library extracted from rat primary cortical cultures, designated here as SGIP1b, conforms to predicted SGIP1 transcript variant X19 (NCBI Reference Sequence: XM_017593774.2). This variant comprises 660 amino acids and differs from the standard SGIP1 variant found on Uniprot (which corresponds to predicted transcript variant X9, NCBI Reference Sequence: XM_017593764.2) by two deletions – a single residue deletion (Q34) in the MP domain and a 165-residue deletion in the PRD (black hatched region). Importantly, SGIP1b does not contain the two additional regions found in SGIP1α, the first of which has been found to be necessary for membrane tubulation (31).The blue cross-hatched region in the µHD indicates the region that binds ctCB1R (23). MP = <u>m</u>embrane <u>p</u>hospholipids binding domain; APA = <u>AP</u>-2 <u>A</u>ctivator motif; PRD = <u>P</u>roline-<u>R</u>ich <u>D</u>omain; µHD = <u>µ H</u>omology <u>D</u>omain. (B) Representative immunoblots showing that FLAG-SGIP1b co-immunoprecipitates with GFP-ctCB1R^WT^, but not GFP-ctCB1R^ΔH9^ or GFP control, in HEK293T cells. (C) Quantification of data represented in (**B**). Significantly more FLAG-SGIP1b co-immunoprecipitates with GFP-ctCB1R^WT^ than with a GFP control (GFP vs. ctCB1R^WT^: mean ± SEM, 4.04 ± 1.95 vs. 100 ± 3.23; ****p < 0.0001) or with GFP-ctCB1R^ΔH9^ (ctCB1R^WT^ vs. ctCB1R^ΔH9^: mean ± SEM, 100 ± 3.23 vs. 8.70 ± 2.38; ***p = 0.0002), suggesting that FLAG-SGIP1b specifically interacts with *H9*. Level of FLAG-SGIP1 co-IP with GFP-ctCB1R^ΔH9^ was comparable to that of the GFP control, suggesting that *H9* is the only interaction site of FLAG-SGIP1b in ctCB1R (GFP vs. ctCB1R^ΔH9^: mean ± SEM, 4.04 ± 1.95 vs. 8.70 ± 2.38; ^ns^p = 0.3340). FLAG-SGIP1 signal was normalised to GFP-IP signal and expressed as a percentage of ctCB1R^WT^. Matched one-way ANOVA with Tukey’s *post hoc* test. n = 5 independent experiments per condition.

Importantly, this variant, which we refer to as SGIP1β, does not contain the additional sequence found in the longer isoform SGIP1α (NCBI Reference sequences NM_001376936.1, transcript variant X1 NCBI Reference Sequence: XM_039109919.1, and X2: NCBI Reference Sequence: XM_039109920.1) that is necessary for membrane tubulation (31). Furthermore, the 99 C-terminal residues D708-N806 in both mouse SGIP1 and rat SGIP1β, which have 100% sequence identity and contain the CB1R binding domain (23), are unchanged.

### SGIP1β interacts with the *H9* domain of CB1R

To determine whether SGIP1 binds CB1R via *H9,* we co-transfected HEK293T cells with FLAG-SGIP1β and either EGFP, EGFP-ctCB1R^WT^, or EGFP-ctCB1R^ΔH9^. Using GFP-Trap, FLAG-SGIP1β co-immunoprecipitated with EGFP-ctCB1R^WT^ but not with EGFP-ctCB1R^ΔH9^ or the EGFP control, indicating that *H9* is required for CB1R binding to SGIP1 (**Figure 1B-C**).

### Overexpression of SGIP1 increases surface expression of CB1R^WT^, but not CB1R^ΔH9^

To determine the role of SGIP1 on CB1R axonal surface expression, we co-transfected DIV12 neurons with full-length EGFP-CB1R^WT^ or EGFP-CB1R^ΔH9^, and either SBP control or an SBP-tagged SGIP1β. Following transfection, neurons were incubated for a further 2 days and then stained for surface CB1R using anti-GFP antibody (**Figure 2A**). Co-expression of SBP-SGIP1β, but not SBP control, significantly increased axonal surface levels of EGFP-CB1R^WT^ (**Figure 2B**), similar to what occurs in HEK293 cells (23). Importantly, no such increase was observed for EGFP-CB1R^ΔH9^, suggesting that *H9* is necessary for this effect to occur (**Figure 2B**).

**Figure 2:**
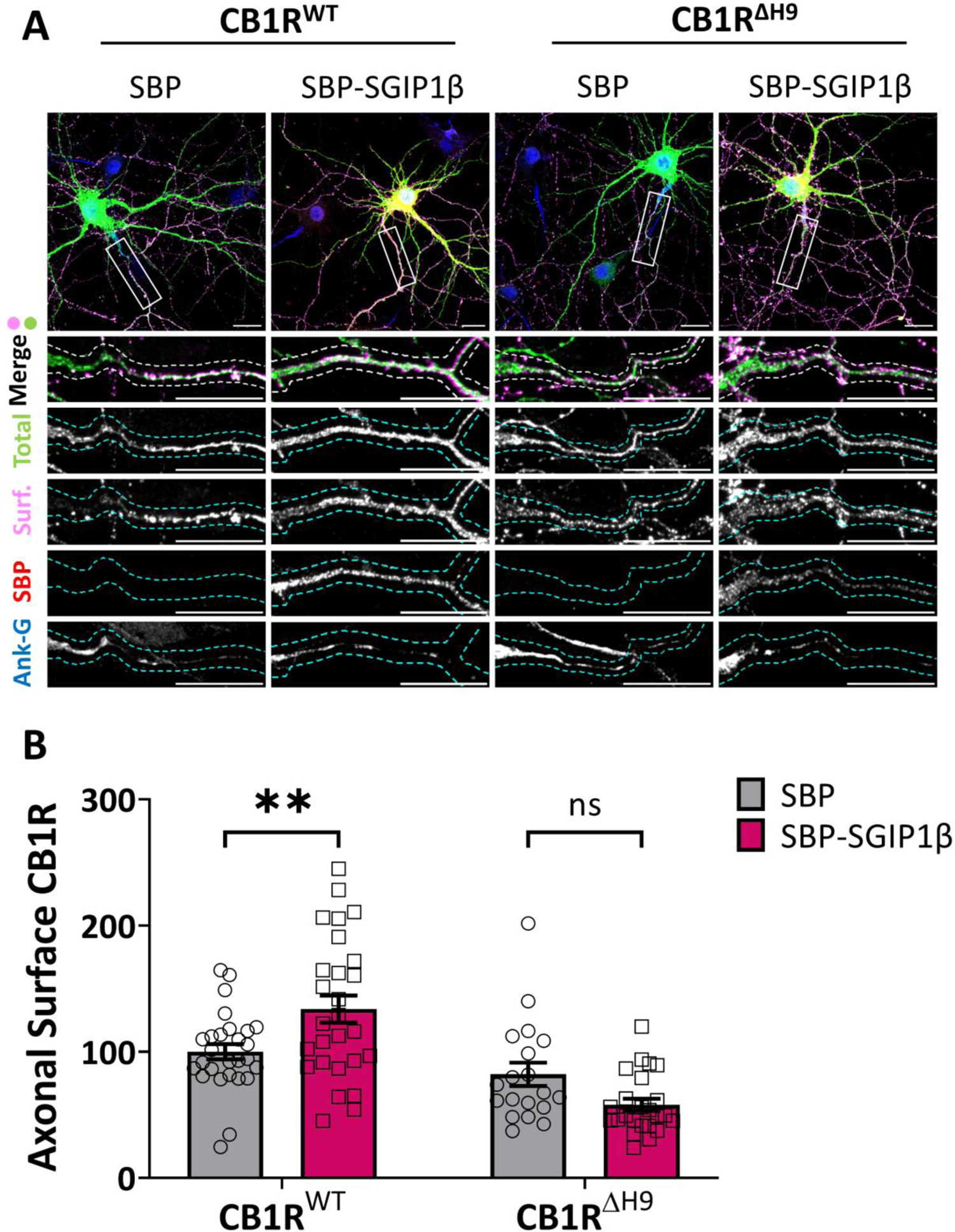
Overexpression of SGIP1β increases surface expression of CB1R^WT^, but not CB1R^ΔH9^. (A) Representative confocal images of DIV14 hippocampal neurons expressing EGFP-CB1R^WT^ or EGFP-CB1R^ΔH9^ and either SBP or SBP-SGIP1b. Cells were transfected at DIV12, incubated for a further 2 days, and then surface stained with anti-GFP antibody. Upper panels for each condition show whole cell field of view and lower panels are enlargements of axonal ROIs. Green = total; magenta = surface; red = SBP-SGIP1; blue = axon marker (Ankyrin-G). Merge: surface to total seen as white. (B) Quantification of data represented in (**A**). SGIP1b overexpression causes a significant increase in surface expression of EGFP-CB1R^WT^ in axons (CB1R^WT^/SBP vs. CB1R^WT^/SBP-SGIP1: mean ± SEM, 100.00 ± 6.03 vs. 133.82 ± 10.72; n = 27 vs. n = 27; **p = 0.0049). SGIP1b overexpression did not alter surface expression of EGFP-CB1R^ΔH9^ (CB1R^ΔH9^/SBP vs. CB1R^ΔH9^/SBP-SGIP1: mean ± SEM, 82.12 ± 9.25 vs. 57.91 ± 4.82; n = 19 vs. n = 24; ^ns^p = 0.100). Surface fluorescence was normalised to total fluorescence and shown as a percentage of CB1R^WT^/SBP control. Two-way ANOVA with Sidak’s *post hoc* test; n = 19-27 neurons per condition from four independent neuronal cultures.

### SGIP1 knockdown reduces surface expression of CB1R^WT^, but not CB1R^ΔH9^

Next, we transfected DIV9 hippocampal neurons with a scrambled shRNA (SCR29), or an shRNA knockdown construct that targets all known isoforms of SGIP1 (25), and either EGFP-CB1R^WT^ or EGFP-CB1R^ΔH9^ (**Supplementary Figure 1**). Following transfection, neurons were incubated for a further 5 days to ensure complete knockdown, and then live stained for surface CB1R using anti-GFP antibody (**Figure 3A**). Consistent with a role for SGIP1 in promoting CB1R axonal surface expression, SGIP1 knockdown reduced surface EGFP-CB1R^WT^ in axons to levels equivalent to EGFP-CB1R^ΔH9^ (**Figure 3B**). Importantly, SGIP1 knockdown did not further reduce surface expression of EGFP-CB1R^ΔH9^ (**Figure 3B**). These data demonstrate that CB1R^ΔH9^ is insensitive to regulation by SGIP1 and strongly suggest that the reduced surface expression phenotype of EGFP-CB1R^ΔH9^ is due to its inability to bind SGIP1.

**Figure 3:**
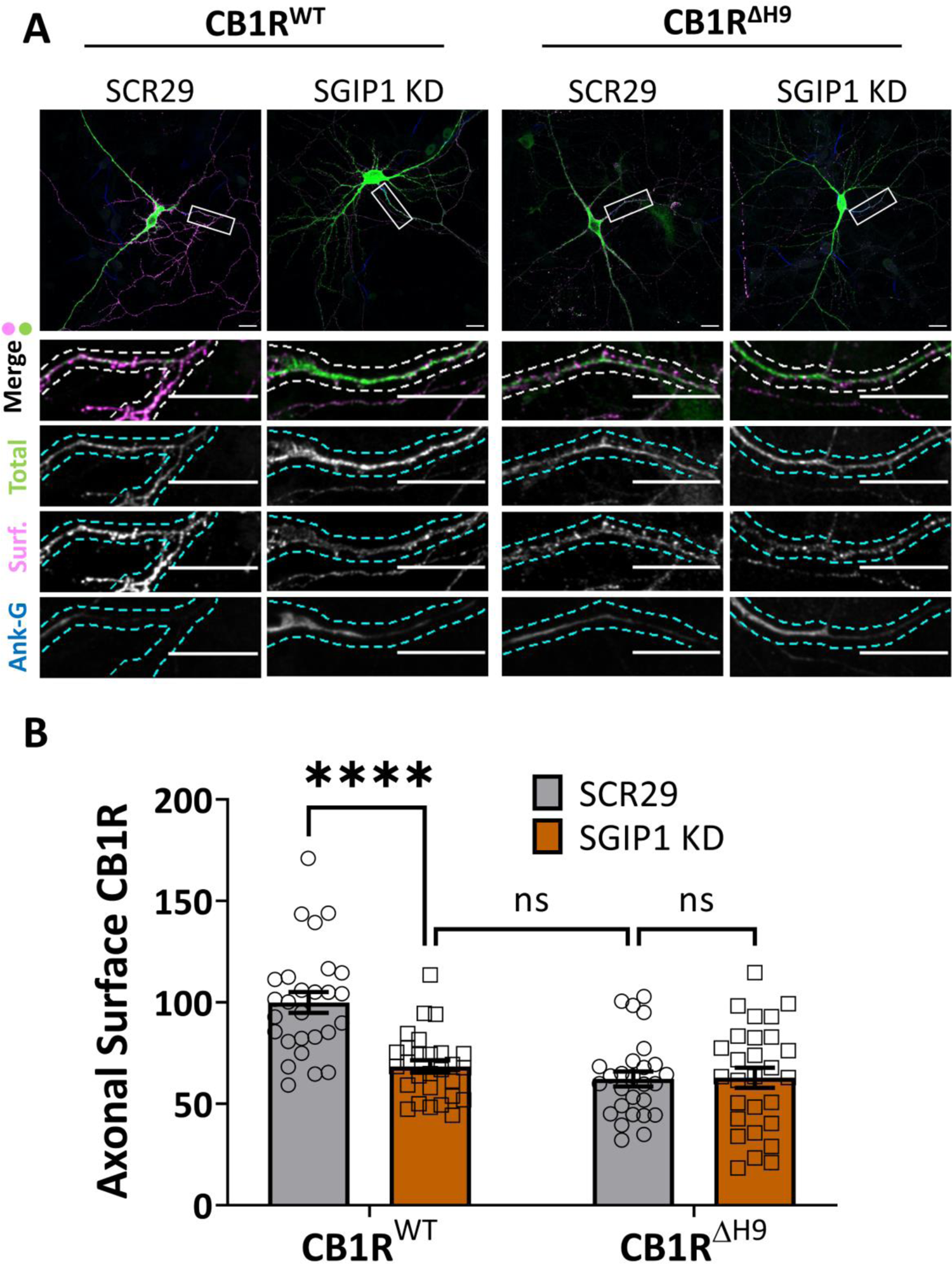
SGIP1 knockdown reduces surface expression of CB1R^WT^, but not CB1R^ΔH9^. (A) Representative confocal images of DIV14 hippocampal neurons expressing EGFP-CB1R^WT^ or EGFP-CB1R^ΔH9^ and either a 25mer shRNA targeting SGIP1 (SGIP1 KD) or a non-targeting scrambled 29mer control (SCR29). Cells were transfected at DIV9 and left for 5 days to ensure knockdown then surface stained with anti-GFP antibody. Upper panels for each condition show whole cell field of view and lower panels are enlargements of axonal ROIs. Green = total; magenta = surface; blue = axon marker (Ankyrin-G). Merge: surface to total seen as white. (B) Quantification of data represented in (**A**). SGIP1 knockdown causes a significant reduction in axonal surface expression of EGFP-CB1R^WT^ (CB1R^WT^/SCR29 vs. CB1R^WT^/SGIP1 KD: mean ± SEM, 100 ± 5.15 vs. 68.46 ± 3.04; n = 27 vs. n = 28; ****p < 0.0001), which phenocopies the reduced surface expression phenotype of EGFP-CB1R^ΔH9^ (CB1R^WT^/SGIP1 KD vs. CB1R^ΔH9^/SCR29: mean ± SEM, 68.46 ± 3.04 vs. 62.31 ± 3.73; n = 28 vs. n = 27; ^ns^p = 0.896). The effect of SGIP1 KD is occluded for EGFP-CB1R^ΔH9^ (CB1R^ΔH9^/SCR29 vs. CB1R^ΔH9^/SGIP1: mean ± SEM, 62.31 ± 3.73 vs. 62.88 ± 4.93; n = 27 vs. n = 28; ^ns^p > 0.999). Surface fluorescence was normalised to total fluorescence and shown as a percentage of CB1R^WT^/SCR29. Two-way ANOVA with Sidak’s *post hoc* test. N = 27-28 neurons from five independent neuronal cultures per condition.

We, and others, have shown that although CB1R is delivered to dendritic plasma membrane, it is rapidly internalised (6, 7, 11, 13, 34). While SGIP1 has been reported to preferentially localise to axons and presynaptic terminals (22, 23, 35), SBP-SGIP1β appears to be present throughout the neuron. However, while no effect of SBP-SGIP1β overexpression on CB1R surface expression was observed in dendrites (**Supplementary Figure 2A**), pan-SGIP1 knockdown reduced dendritic surface levels of CB1R^WT^, but not CB1R^ΔH9^ (**Supplementary Figure 2B**). These results raise the possibility that an isoform other than SGIP1β might affect CB1R surface expression in dendrites.

### SGIP1 knockdown reduces the surface/total ratio of endogenous CB1R

To determine how SGIP1 effects surface expression of endogenous CB1R, we transduced DIV7/8 primary cortical neurons with lentivirus expressing either a scrambled shRNA control (SCR29) or SGIP1 shRNA. Surface and total levels of endogenous CB1R were examined at DIV14/15 by surface biotinylation followed by streptavidin pulldown and Western blotting (**Figure 4A**). Consistent with our exogenously expressed CB1R data, knockdown of SGIP1 to levels of ∼15% compared to the SCR29 control significantly decreased surface expression of endogenous CB1R (**Figure 4B-C** and **Supplementary Figure 1**). These results further support a role for SGIP1 in promoting CB1R surface expression.

**Figure 4:**
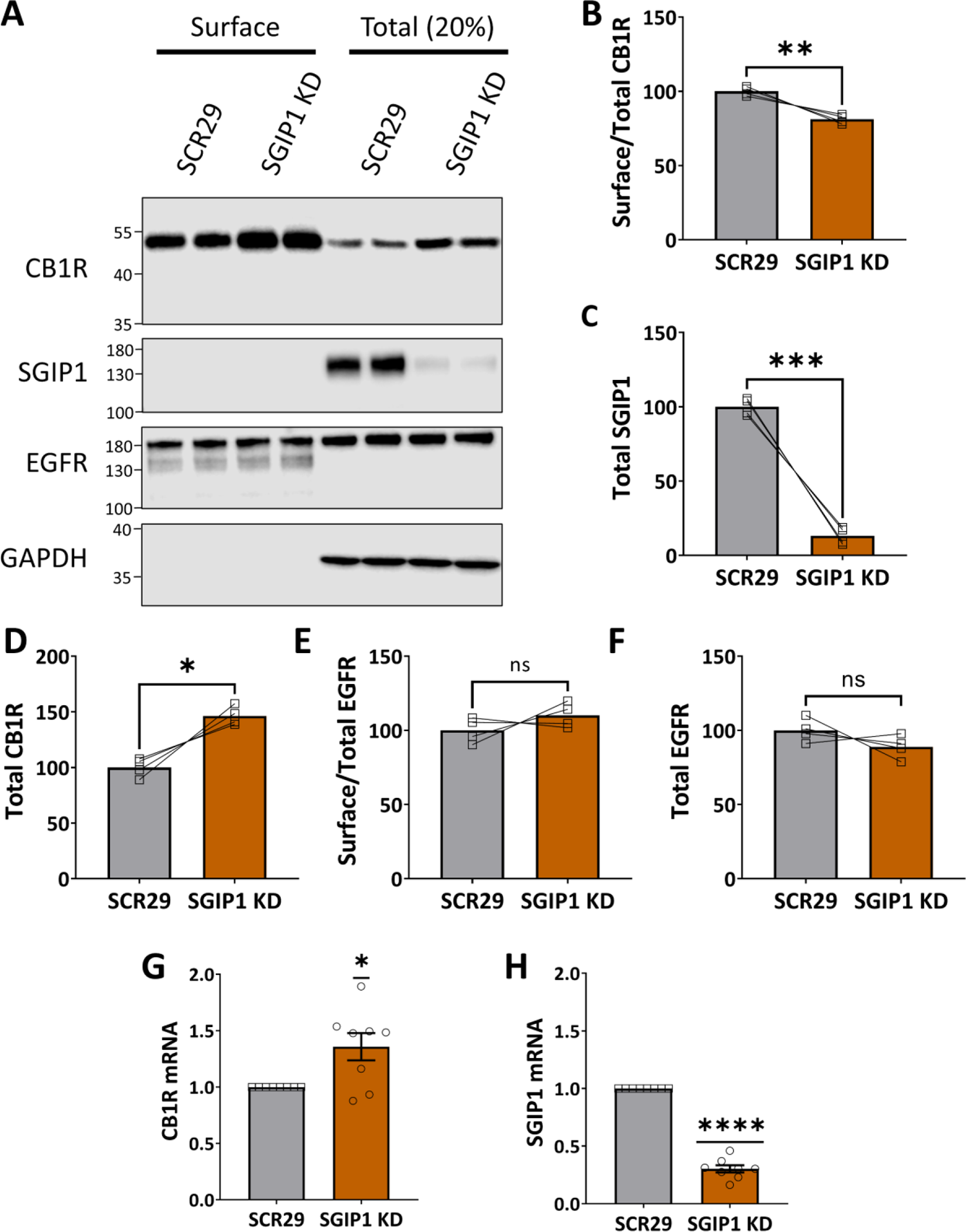
SGIP1 knockdown decreases the surface/total ratio of endogenous CB1R. (A) Representative immunoblots showing surface (left) and total (right; 20%) levels of endogenous CB1R in DIV14 cortical neurons transduced with SCR29 or SGIP1 shRNA. Blots were probed with anti-SGIP1 to determine KD efficiency. Only experiments with >80% KD efficiency of SGIP1 KD were analysed. EGFR was included as a control surface protein and GAPDH as a loading control. (**B-F**) Quantification of data represented in (**A**), expressed as percentage of SCR29 control. Paired t-tests; n = four independent experiments (each experiment = average of duplicates). (B) Surface/total ratio of CB1R is significantly reduced with SGIP1 KD compared to SCR29 control (SCR29 vs. SGIP1 KD: mean ± SEM, 100.00 ± 1.48 vs. 81.23 ± 1.48, t(3) = 6.364, **p = 0.0079). (**C**) Total levels of CB1R were significantly increased with SGIP1 KD compared to SCR29 control (SCR29 vs. SGIP1 KD: mean ± SEM, 100.00 ± 4.22 vs. 146.40 ± 4.22, t(3) = 5.493, *p = 0.0119). (**D**) SGIP1 levels were knocked down by about 87% compared to SCR29 control (SCR29 vs. SGIP1 KD: mean ± SEM, 100.00 ± 2.82 vs. 13.10 ± 2.82, t(3) = 15.42, ***p = 0.0006). (**E**) SGIP1 KD had no significant effect on EGFR surface/total levels (SCR29 vs. SGIP1 KD: mean ± SEM, 100.00 ± 4.18 vs. 110.10 ± 4.18, t(3) = 1.210, ^ns^p = 0.138). (**F**) SGIP1 KD had no significant effect on EGFR total levels (SCR29 vs. SGIP1 KD: mean ± SEM, 100.00 ± 3.94 vs. 81.23 ± 3.94, t(3) = 1.406, ^ns^p = 0.094). (**G-H**) RT-qPCR of DIV14/15 cortical neurons transduced with SCR29 or SGIP1 KD lentivirus. Cycle threshold (Ct) values were normalised to GAPDH (ΔCt) and presented as fold change compared to SCR29 control 2^(-ΔΔCt)^. One sample t-tests (theoretical mean = 1); n = eight independent experiments. (**G**) KD of SGIP1 increases CB1R mRNA levels (SGIP1 KD: mean ± SEM, 1.367 ± 0.1206 vs. theoretical mean = 1.000, n = 8, t(7) = 2.963, *p = 0.0210). (**H**) SGIP1 KD lentivirus reduced SGIP1 transcript levels by about 70% (SGIP1 KD: mean ± SEM, 0.3027 ± 0.03127 vs. theoretical mean = 1.000, n = 8, t(7) = 22.30, ****p < 0.0001).

Surprisingly, however, in these experiments, total levels of endogenous CB1R were increased by SGIP1 knockdown compared to the SCR29 control (**Figure 4D**). In contrast, neither total (**Figure 4E**) nor surface (**Figure 4F**) levels of epidermal growth factor receptor (EGFR) were affected by SGIP1 knockdown, suggesting the changes observed could not be attributed to global alterations in membrane protein expression or surface expression.

Therefore, to assess if the increase in total CB1R protein levels could be due to increased transcription, we analysed transcript levels of CB1R in DIV14/15 cortical cells transduced with SCR29 or SGIP1 shRNA by RT-qPCR (**Figure 4G-H** and **Supplementary Figure 3**). The relative mRNA level of CB1R was significantly increased when SGIP1 was knocked down, suggesting that enhanced CB1R levels likely constitute a homeostatic mechanism in response to reduced surface expression.

### SGIP1 knockdown impairs CB1R-mediated inhibition of intracellular Ca^2+^ influx

We next investigated how presynaptic CB1R signalling is affected by loss of SGIP1. CB1R/SGIP1 co-expression in cell lines alters CB1R-mediated ERK1/2 phosphorylation as well as recruitment of β-arrestin and GRK3 (23, 32), whereas G protein activation and intracellular Ca^2+^ mobilisation is unaffected (23).

To define what happens in neurons we transfected DIV8/9 primary hippocampal neurons with the presynaptically-localised Ca^2+^ indicator synaptophysin-GCaMP3 (SyGCaMP3) (36) and either SCR29 or SGIP1 knockdown shRNA, and assayed CB1R function at DIV14/15. Neurons were stimulated with 20 APs (50V, 1ms pulses) at 20Hz, perfused with the CB1R agonist 2-AG (1 μM) for 3 minutes, and restimulated. The Ca^2+^ signal was compared before and after 2-AG incubation (**Figure 5A**).

**Figure 5:**
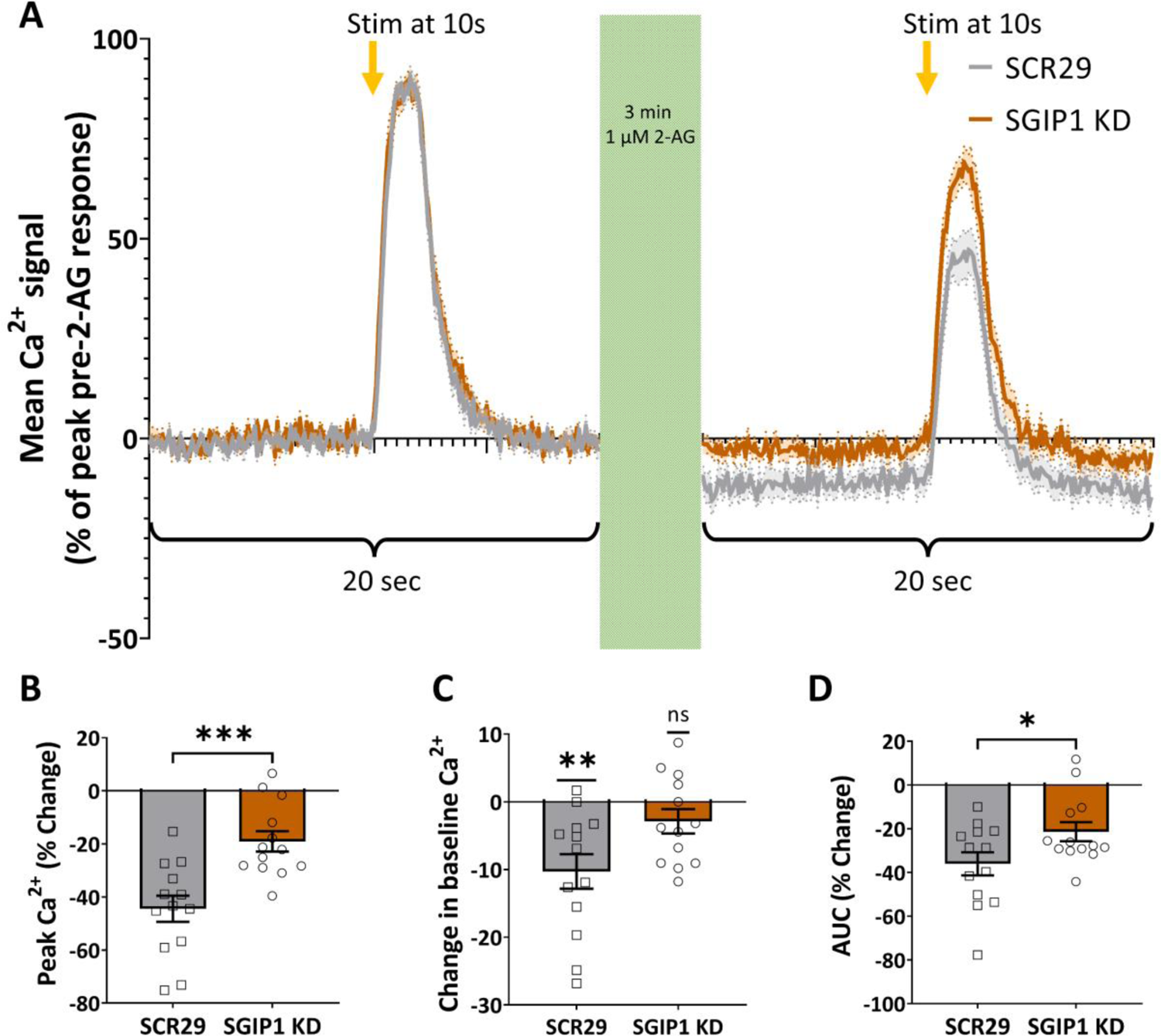
SGIP1 knockdown decreases CB1R-mediated inhibition of Ca^2+^ influx. (**A**) Average traces of presynaptic Ca^2+^ responses (measured using SyGCaMP3) to 20 action potentials at 20 Hz pre- and post-2-AG incubation (1 μM, 3 min). Data is expressed as a percentage of pre-2-AG peak. Solid line = mean; shading and dotted line = SEM; n = 13 fields of view from eight independent neuronal cultures. (**B-D**) Quantification of data represented in (**A**). (**B**) SGIP1 KD significantly reduces the drop in peak Ca^2+^ levels after 2-AG application compared to SCR29 control (SCR29 vs. SGIP1 KD: mean ± SEM, −45.45 ± 4.92 vs. −19.07 ± 3.84; n = 13 vs. n = 13, t(24) = 4.066, ***p = 0.0004). Unpaired t-test. n = 13 fields of view from eight independent neuronal cultures. (**C**) 2-AG application significantly reduces the baseline level of Ca^2+^ of SCR29-transfected control neurons, but not of SGIP1 KD neurons (SCR29 vs. 0: mean ± SEM, −10.29 ± 2.55; n = 13, t(12) = 0.0017, **p = 0.0017. SGIP1 KD vs. 0: mean ± SEM, −2.89 ± 1.80; n = 13, t(12) = 1.607, ^ns^p = 0.134). One sample t-test with theoretical mean = 0; n = 13 fields of view from eight independent neuronal cultures. (**D**) SGIP1 KD significantly reduces the area under curve (AUC) of Ca^2+^ signal after 2-AG application compared to SCR29 control (SCR29 vs. SGIP1 KD: mean ± SEM, −36.06 ± 5.30 vs. −21.36 ± 4.38; n = 13 vs. n = 13, t(24) = 2.139, *p = 0.043). Unpaired t-test; n = 13 fields of view from eight independent neuronal cultures.

As expected, in control neurons the peak Ca^2+^ signal decreased after 2-AG incubation by ∼45% (**Figure 5B**), consistent with the presynaptic inhibitory action of CB1R signalling (37, 38). The magnitude of this decrease was markedly reduced in SGIP1 knockdown neurons (**Figure 5B**). Furthermore, 2-AG significantly decreased the baseline Ca^2+^ signal compared to pre-2-AG incubation in control cells, but not after SGIP1 knockdown (**Figure 5C**). Lastly, the reduction in the area under the curve of the Ca^2+^ response was significantly less in SGIP1 knockdown cells compared to control cells (**Figure 5D**). Together, these data suggest that SGIP1 knockdown suppresses the endocannabinoid-mediated reduction in Ca^2+^ influx, indicative of reduced presynaptic CB1R signalling in the absence of SGIP1.

## Discussion

The context of this study was that the amphipathic α-helical *H9* domain in the intracellular C-terminal of CB1R contributes to the polarised presynaptic surface expression of CB1R (13). SGIP1 binds to CB1R, increasing its surface expression and modulating its signalling in HEK293 cells (23, 32). The distributions of SGIP1 and CB1R overlap in mouse brain (39), and they co-localise at the presynapse (23). We therefore hypothesised that SGIP1 might interact with *H9* to modulate synaptic CB1R availability at the presynaptic membrane.

We show that SGIP1 binds to the CB1R *H9* domain to promote axonal surface expression. A FLAG-tagged isoform of SGIP1, which we refer to as SGIP1β, co-IPs with ctCB1R^WT^, but not with ctCB1R^ΔH9^ in HEK293T cells (**Figure 1A-C**). Moreover, overexpression of SGIP1β increases CB1R axonal plasma membrane expression (**Figure 2**), while knockdown of SGIP1 reduces axonal surface levels (**Figure 3**). Importantly, neither overexpression (**Figure 2**) nor knockdown (**Figure 3**) of SGIP1 altered CB1R^ΔH9^ surface expression, consistent with SGIP1 mediating its effect through interaction with the *H9* domain of CB1R.

SGIP1 knockdown reduces both the surface/total ratio of endogenous CB1R (**Figure 4A-C**) and modulates downstream CB1R signalling. Under control conditions, CB1R activation inhibits voltage-gated Ca^2+^ channels via G_i_ βγ-subunit mobilisation (37, 38), but the magnitude of the effect is significantly reduced with SGIP1 knockdown (**Figure 5**). Notably, this effect on signalling remains even though total CB1R levels increase with SGIP1 knockdown (**Figure 4D**) – a phenomenon that we speculate may represent a homeostatic mechanism in response to decreased CB1R surface expression, since CB1R transcript levels were also increased (**Figure 4G-H**). Taken together, these data provide strong evidence that SGIP1 promotes axonal surface localisation of CB1R through interaction with *H9*.

SGIP1 is a member of the muniscin family of cargo adapters due to its similarity with FCHo1/2 proteins and interacts with AP-2 (40), intersectin (28), and Eps15 (25). Since SGIP1 is a component of the clathrin-mediated endocytosis complex, a key question is why does SGIP1 overexpression *enhance*, and knockdown *reduce,* CB1R surface expression?

We speculate that key to untangling this conundrum is the observation that the actions of SGIP1 are highly isoform-dependent (31). Both SGIP1 and FCHo1/2 proteins contain an N-terminal membrane phospholipid (MP) binding domain, an AP-2 activator domain, a proline rich domain, and a C-terminal µ homology domain (28, 40) (**Figure 1A**). However, the membrane binding domain of FCHo1/2 is an F-BAR domain that deforms the plasma membrane to facilitate clathrin-coated pit formation. The corresponding region of SGIP1, however, has no F-BAR sequence similarity.

Nonetheless, a recent report has identified a 28-residue, positively charged sequence present in the membrane binding domain that is necessary for homo-oligomerisation of SGIP1 and membrane tubulation (31). Of the 32 different predicted rat transcript variants available on the NCBI database, 13 contain the membrane tubulating sequence in its entirety. Importantly, SGIP1α contains this sequence, while SGIP1 and SGIP1β do not, and previous experiments indicating that SGIP1/CB1R co-expression in HEK293 cells increases CB1R surface levels were performed using the non-membrane tubulating form of SGIP1 (23). Consistent with these findings, our data show that overexpression of SGIP1β, which does not contain the tubulating sequence, increases CB1R surface levels in axons (**Figure 2**).

We note, however, that different SGIP1 isoforms may be selectively recruited to different cargo to mediate opposing effects. Differences between the SGIP1 isoforms are in the homo-oligomerisation and membrane-tubulating sequences, whereas the C-terminal 99 residue domain of SGIP1 that interacts with CB1R is present in all three SGIP1 isoforms. One possibility could be that SGIP1 and SGIP1β act as endogenous ‘dominant negatives’ to SGIP1α and FCHo1/2 proteins, preventing them from binding cargo by taking up the binding site. However, further work will be required examine this possibility directly. In conclusion, our findings indicate that SGIP1 promotes CB1R surface expression via interaction with *H9*.

## Materials and methods

### Plasmids and reagents

To N-terminally tag rat CB1R, the first 25 N-terminal amino acids were removed and an exogenous signal peptide corresponding to interleukin-2 (SP^Il2^) was added before the tags as previously characterised (8, 13).

The following overexpression plasmids were used:

| Vector | Overexpression | Figure |
| --- | --- | --- |
| pEGFP-C2 | EGFP | 1 |
| pEGFP-C2 | EGFP-ctCB1R<sup>WT</sup> | 1 |
| pEGFP-C2 | EGFP-ctCB1R<sup>ΔH9</sup> | 1 |
| pcDNA3.1 | FLAG-SGIP1β | 1; S2 |

The following double overexpression plasmids (one cassette with a CMV promoter, one cassette with an Sffv promotor) were used:

| Vector | CMV- | Sffv- | Figure |
| --- | --- | --- | --- |
| pXlg3 | SBP | SP<sup>IL2</sup>-EGFP-CB1R<sup>WT</sup> | 2 |
| pXlg3 | SBP-SGIP1 | SP<sup>IL2</sup>-EGFP-CB1R<sup>WT</sup> | 2 |
| pXlg3 | SBP | SP<sup>IL2</sup>-EGFP-CB1R<sup>ΔH9</sup> | 2 |
| pXlg3 | SBP-SGIP1 | SP<sup>IL2</sup>-EGFP-CB1R<sup>ΔH9</sup> | 2 |

The following shRNA knockdown (H1 promoter) + overexpression (Sffv promoter) plasmids were used:

| Vector | shRNA KD | Overexpression | Figure |
| --- | --- | --- | --- |
| pXlg3 | SCR29 | SP<sup>IL2</sup>-EGFP-CB1R<sup>WT</sup> | 3 |
| pXlg3 | SGIP1 KD | SP<sup>IL2</sup>-EGFP-CB1R<sup>WT</sup> | 3 |
| pXlg3 | SCR29 | SP<sup>IL2</sup>-EGFP-CB1R<sup>ΔH9</sup> | 3 |
| pXlg3 | SGIP1 KD | SP<sup>IL2</sup>-EGFP-CB1R<sup>ΔH9</sup> | 3 |
| pSUPER.neo | SCR29 | mCherry | S2 |
| pSUPER.neo | SGIP1 KD | mCherry | S2 |
| pXlg3 | SCR29 | mCherry | 4; S2 |
| pXlg3 | SGIP1 KD | mCherry | 4; S2 |
| pXlg3 | SCR29 | SyGCaMP3 | 5 |
| pXlg3 | SGIP1 KD | SyGCaMP3 | 5 |

| **Antibody** | **Clone** | **Source** | **Identifier** | **Concentration** |
| --- | --- | --- | --- | --- |
| anti-GFP<br>(chicken) | Poly | Abcam | Cat# ab13970,<br>RRID:AB_300798 | ICC: 1:1,000 |
| Anti-Ankyrin G<br>(mouse) | N106/<br>36 | UC Davis/NIH<br>NeuroMab<br>Facility | Cat# N106/36,<br>RRID:AB_2877524 | ICC: 1:500 |
| Anti-Ankyrin G<br>(rabbit) | Poly | Synaptic Systems | Cat# 386 003,<br>RRID:AB_2661876 | ICC: 1:500 |
| Anti-SBP-tag<br>(mouse) | 20 | Millipore | Cat# MAB10764,<br>RRID:AB_10631872 | ICC: 1:500 |
| Anti-FLAG<br>(mouse) | M2 | Sigma-Aldrich | Cat# F1804,<br>RRID:AB_262044 | WB: 1:2,000-1:10,000 |
| Anti-GFP<br>(rat) | 3H9 | ChromoTek | Cat# 3h9-100,<br>RRID:AB_10773374 | WB: 1:2,000-1:5,000 |
| Anti-CB1R<br>(rabbit) | Poly | Abcam | Cat# ab23703,<br>RRID:AB_447623 | WB: 1:250-1:1,000 |
| Anti-SGIP1<br>(rabbit) | Poly | Sigma-Aldrich | Cat# HPA017963,<br>RRID:AB_10964882 | WB: 1:100 |
| Anti-SGIP1<br>(rabbit) | Poly | Biorbyt | Cat# orb40255,<br>RRID:AB_10997021 | WB: 100 |
| Anti-EGFR | EP38Y | Abcam | Cat# ab52894,<br>RRID:AB_869579 | WB: 1:1,000 |
| Anti-GAPDH | 6C5 | Abcam | Cat# ab8245,<br>RRID:AB_2107448 | WB: 1:10,000-1:40,000 |

The non-targeting 29-mer shRNA (SCR29) sequence was:

5’ – GCACTACCAGAGCTAACTCAGATAGTACT – 3’ (Origene)

The SGIP1 shRNA target sequence was:

5’ – CCAATACCAAGGAATTCTGGGTAAA – 3’ (25).

The lentiviral helper vectors p8.91 and pMD2.G were used. The following primary antibodies were used:

The following secondary antibodies were used:

| **Antibody** | **Conjugate** | **Source** | **Identifier** |
| --- | --- | --- | --- |
| Anti-chicken IgY<br>(donkey) | Alexa Fluor<br>647 | Jackson<br>ImmunoResearch Labs | Cat# 703-606-155,<br>RRID:AB_2340380 |
| Anti-chicken IgY<br>(donkey) | Cy2 | Jackson<br>ImmunoResearch Labs | Cat# 703-225-155,<br>RRID:AB_2340370 |
| Anti-mouse IgG<br>(donkey) | Cy3 | Jackson<br>ImmunoResearch Labs | Cat# 715-165-150,<br>RRID:AB_2340813 |
| Anti-rabbit IgG<br>(goat) | DyLight 405 | Jackson<br>ImmunoResearch Labs | Cat# 111-475-003,<br>RRID:AB_2338035 |
| Anti-mouse IgG<br>(goat) | DyLight 405 | Jackson<br>ImmunoResearch Labs | Cat# 115-475-003,<br>RRID:AB_2338786 |
| Anti-mouse IgG<br>(goat) | HRP | Sigma-Aldrich | Cat# A3682,<br>RRID:AB_258100 |
| Anti-rabbit IgG<br>(goat) | HRP | Sigma-Aldrich | Cat# A6154,<br>RRID:AB_258284 |
| Anti-rat IgG (goat) | HRP | Sigma-Aldrich | Cat# A5795,<br>RRID:AB_258259 |

All fluorescent secondary antibodies for ICC were used at 1:400 and all HRP secondary antibodies for WB were used at 1:10,000.

### Cell culture and transfection

The hippocampus and cortex were dissected from E17 Han Wistar rats and dissociated according to standard protocols (41). 200,000-350,000 dissociated hippocampal neurons were plated onto 25 mm glass coverslips pre-coated with PDL (Sigma) in plating media (Neurobasal, Gibco; supplemented with 5% horse serum, Sigma; 2 mM GlutaMAX, Gibco; and 1xGS21, GlobalStem). 500,000 cortical neurons were plated per well of a 6 well plate pre-coated with PDL. The next day, plating media was removed and replaced with feeding media (Neurobasal supplemented with 1.2 mM GlutaMAX and 1xGS21). When GS21 became unavailable, dissociated hippocampal neurons were both plated and maintained in B27 Plus media (Neurobasal Plus supplemented with 1x B27 Plus, Gibco).

DIV9 (KD) or DIV12 (overexpression) primary hippocampal neurons were transfected with 1-2 μg plasmid DNA and 5 μL Lipofectamine 2000 (Invitrogen) according to manufacturer’s instructions, with minor modifications. For overexpression experiments, cells were left 2-3 days whereas for KD experiments, cells were left 5 days to ensure efficient KD. All experiments were carried out at DIV14/15. Cortical neurons were transduced with lentivirus at DIV7/8 and assayed at DIV14/15. To limit glial growth, antimitotics (FDU + uridine, final concentration 0.4 μM) were added at DIV7/8 to cortical cultures.

HEK293T cells (EACC) were passaged and maintained in glutamine-containing DMEM (Gibco) supplemented with 10% FBS and 1% penicillin/streptomycin and were transfected with Lipofectamine 2000 (Invitrogen) in penicillin/streptomycin-free media for 48 hours according to manufacturer’s instructions, with minor modifications. HEK293T cells were treated with ciprofloxacin (10 μg/mL) regularly to prevent mycoplasma contamination.

### GFP-Trap

GFP was immunoprecipitated with GFP-Trap (Chromotek) according to manufacturer’s instructions, with minor modifications. Samples were kept at 4°C throughout. HEK293T cells were lysed in lysis buffer (50 mM Tris HCl pH 7.4, 150 mM NaCl, 0.5% Triton X, 1 x cOmplete protease inhibitors [1x = 1 tablet in 40mL; Merck]), sonicated, incubated 20 min, and clarified by centrifugation at 16k RCF for 20 min. A proportion of the lysate was kept aside (“input”), and the rest was incubated with GFP-Trap beads on a rotating wheel for 1 hour. The beads were pelleted for 2 minutes at 1.5k RCF and washed 3x in wash buffer (lysis buffer minus protease inhibitors). 2x Laemmli sample buffer was added to the beads and the inputs and the beads were boiled at 95°C for 5 minutes.

### Lentivirus production and transduction of cortical neurons

Lentivirus was produced in HEK293T cells following standard protocols (42). HEK293T cells plated in 6- or 10-cm dishes were transfected using plain DMEM containing 2 μg/mL of the appropriate pXlg3 viral vector, 0.5 μg/mL pMD2.G, 1.5 μg/mL p8.91, and 12 μg/mL PEI in plain DMEM media for 4 hours. The transfection mix was removed and replaced with DMEM + 10% FBS. After 48 hours, the virus-containing media was collected, centrifuged at 4k rpm for 10 min, and passed through a 0.45 μm syringe filter to remove any remaining HEK293T cells. 500 μL of virus was added per well of a 6 well plate of DIV7 cortical neurons in duplicate and incubated for 7 days.

### Surface biotinylation and streptavidin pulldown

All solutions were pre-chilled to 4°C and steps were carried out on ice. DIV14/15 cortical neurons in 6-well plates transduced with lentivirus were cooled on ice to prevent endocytosis, then washed 3x in ice-cold 1xPBS. Surface proteins were biotinylated by incubation with 0.3 mg/mL of EZ-link Sulfo-NHS-SS-Biotin (Thermo) dissolved in 1xPBS for 10 minutes. Unreacted biotin was washed off with 3x washes in PBS and quenched with a 2 min incubation in 50 mM NH_4_Cl in PBS. Quenching solution was washed off with an additional 3 washes in PBS and cells were lysed in 250 μL of lysis buffer (50 mM Tris HCl pH 7.4, 150 mM NaCl, 1% CHAPS, 0.1% SDS, 10% glycerol, 1 mM EDTA, 1 x cOmplete protease inhibitors), sonicated, incubated 20 min, and clarified by centrifugation at 16k RCF for 20 min.

Biotinylated surface proteins were isolated using streptavidin-coated agarose beads (Merck). The beads were washed 2 times in lysis buffer by centrifugation at low speed (<1.5k RCF). 50 μL of clarified lysate was set aside (total) and 100 µL of clarified lysate was added to 30 µL of beads along with 500 µL of lysis buffer (surface). The beads were incubated on a rotating wheel at 4°C for 1.5 hours, then washed in wash buffer (lysis buffer without protease inhibitors) 3 times, by pelleting the beads for 2 minutes at 1k RCF and discarding the supernatant between washes. 2x Laemmli sample buffer was added to both the surface and total samples. The samples were vortexed, spun down, and incubated overnight at RT (to prevent CB1R aggregation).

### Western Blotting

Samples in Laemmli sample buffer were resolved on 10% acrylamide gels by SDS-PAGE, transferred onto methanol-activated PVDF membrane, and immunoblotted according to standard protocols. Chemiluminescence was detected using a LI-COR Odyssey Fc and quantified using Image Studio.

### RT-qPCR

RNA was extracted from DIV14/15 cultured cortical cells transduced with lentivirus expressing a 29-mer non-targeting shRNA (SCR29) or SGIP1-targeting shRNA using the Qiagen RNeasy mini kit following manufacturer’s instructions. RNA concentration was measured using a nanodrop and 1 µg of RNA was converted to cDNA using the RevertAid First Strand cDNA Synthesis Kit (ThermoFisher Scientific) according to manufacturer’s instructions.

qPCR was performed using PowerUp™ SYBR™ Green Master Mix (ThermoFisher Scientific) mixed with 2 µL of each sample and gene-specific primers at 0.25 µM each and run on a qPCR machine with MxPro software and SYBR green with dissociation curve setup.

The following primers were used:

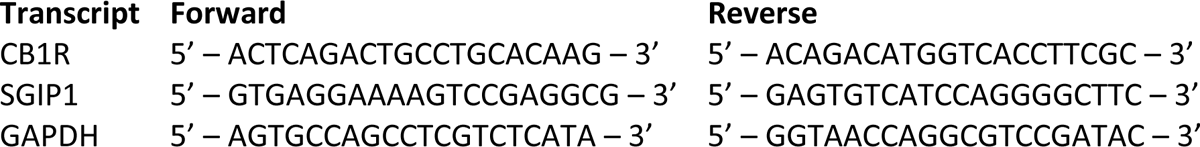

Unknown samples were run in triplicate and no-RT (-RT), no template (NTC), and SYBR neg (10 µL SYBR master mix + 10 µL H_2_O) controls were included with each qPCR.

Relative gene expression was quantified using the ΔΔCt method. Mean cycle threshold (Ct) values of the gene of interest were normalised first to GAPDH mean Ct values (ΔCt) and then to the SCR29 control (ΔΔCt). Fold change of gene expression was plotted as 2^-ΔΔCt^ and a one sample t-test was performed to determine whether the SGIP1 KD condition was significantly different from 1.

### Live surface staining

To measure surface expression, DIV14 cultured hippocampal neurons grown on 25mm glass coverslips were incubated live in the appropriate antibody raised against an extracellular epitope. Briefly, cells were removed from the incubator and allowed to cool to RT for 5 minutes. Cells were then incubated in chicken anti-GFP antibody (1:1,000) in 90 μL conditioned media for 10 minutes at RT. The antibody mix was dotted onto parafilm, and the coverslips were incubated upside down to ensure even coating. Cells were washed 3 times in 1xPBS to remove excess antibody and fixed.

### Fixation and fixed immunostaining

Cells were fixed in pre-warmed 4% PFA and 5% sucrose in 1xPBS for 12 minutes. Following 3 washes in PBS, residual PFA was quenched with a wash in 100mM glycine in 1xPBS, and the cells were washed 3 more times in PBS.

Cells were blocked and permeabilised in 3% BSA in 1xPBS with 0.1% Triton-X for 20 minutes. Cells were then incubated in secondary antibody to label surface. Cells were then stained for total levels and Ankyrin-G (axon initial segment marker). Primary and secondary antibodies were diluted in 3% BSA in 1xPBS. 90 μL of the antibody mix was dotted onto parafilm and the coverslips were incubated upside down for 1 hour at RT. The cells were washed 3x in PBS between incubations. Coverslips were dipped in distilled H_2_0 and mounted onto slides using Fluoromount G (ThermoFisher Scientific) mounting media with or without DAPI.

### Fixed image acquisition and analysis

Images were acquired using a Leica SP8 confocal laser scanning microscope (Wolfson Bioimaging Facility, University of Bristol). All settings were kept the same within experiments. To avoid bias, neurons were selected for data acquisition based only on their total staining and surface was stained in far red.

Fiji (ImageJ) was used to quantify fluorescence. Images were max projected, and regions of interest (ROIs) of approximately similar lengths were drawn around axons based on the total channel only. Axons were defined as processes whose initial segment was positive for Ankyrin-G. Surface fluorescence was normalised to total fluorescence to control for differences in expression levels and expressed as a percentage of the control condition.

### SyGCaMP assay and analysis

SyGCaMP assay was performed as previously described (36). Hippocampal neurons grown on 25mm coverslips were transfected at DIV8/9 with pXlg3-SCR29-SyGCaMP3 or pXlg3-SGIP1 KD-SyGCaMP3 and assayed at DIV14/15. Cells were assayed in HEPES Buffered Saline (HBS; 90-140 mM NaCl [adjusted according to osmolarity of culture media], 5mM KCl, 1.8 mM CaCl_2_, 0.8mM MgCl_2_, 25 mM HEPES pH 7.4, 5 mM glucose) with 25 µM CNQX and 50 µM D-AP5 to prevent spontaneous firing. Timelapse imaging of SyGCaMP was performed at 10Hz with 2×2 binning on a Nikon Eclipse Ti-E C1 plus widefield microscope with a 40x objective, a CCD camera, a GFP filter cube, and accommodating an electrical field stimulation setup. A 20s imaging session was started. After 10s of baseline recording, cells were electrically stimulated with 20 APs (50V, 1ms pulses) at 20Hz. Cells were then perfused with 1 μM 2-AG and incubated for 3 minutes, and the 20s recording and stimulation were repeated.

For each field of view, the mean fluorescence of 3-10 punctate regions of interest (ROIs) were analysed, and non-responsive ROIs were discarded. The mean fluorescence of 3 background ROIs was also measured, which was subtracted for each timepoint for each ROI. Background subtracted values then normalised to basal levels for each ROI (calculated as average mean fluorescence during 8-9s). The average for each timepoint of all ROIs for a field of view was found and then plotted as a percentage of the peak mean fluorescence of the first stimulation. Three parameters were analysed: 1) change in peak post-2-AG signal, 2) change in baseline (calculated as average during 8-9s post-2-AG), 3) change in area under curve (AUC; Total Peak Area was calculated using Prism with the following parameters: baseline = mean of 8-9s and 18-19s, ignoring peaks less than 10% distance from minimum to maximum Y, ignoring any peaks defined by fewer than 8 adjacent points. Post-2-AG AUC was presented as a percentage of pre-2-AG AUC).

### Statistics

All statistics were performed using GraphPad Prism (version 9). Outliers were removed using GraphPad Prism’s ROUT method (Q = 1%). T-tests were used to determine statistical significance between two groups. For more than two groups, One- or Two-way ANOVAs with Tukey’s or Sidak’s *post hoc* tests were used to determine statistical significance, depending on the comparisons required.

For image analysis, ‘n’ denotes the total number of neurons that were analysed, as is convention in the field (6–8, 11, 13). However, the number of separate neuronal cultures prepared from litters of pups from separate dams is also noted for each experiment. For surface biotinylation experiments, ‘n’ denotes the number of separate neuronal cultures, where each ‘n’ is the average of two duplicate experiments. For all data, *p ≤ 0.05, **p ≤ 0.01, ***p ≤ 0.001, ****p ≤ 0.0001. Data are presented as mean ± SEM.

## Acknowledgements

We are grateful to the Wolfson Bioimaging Facility (University of Bristol) and to the Wellcome Trust (220799/Z/20/Z) and the BBSRC (BB/R00787X/1) for financial support.

## Supplementary Figures

**Supplementary Figure 1:**
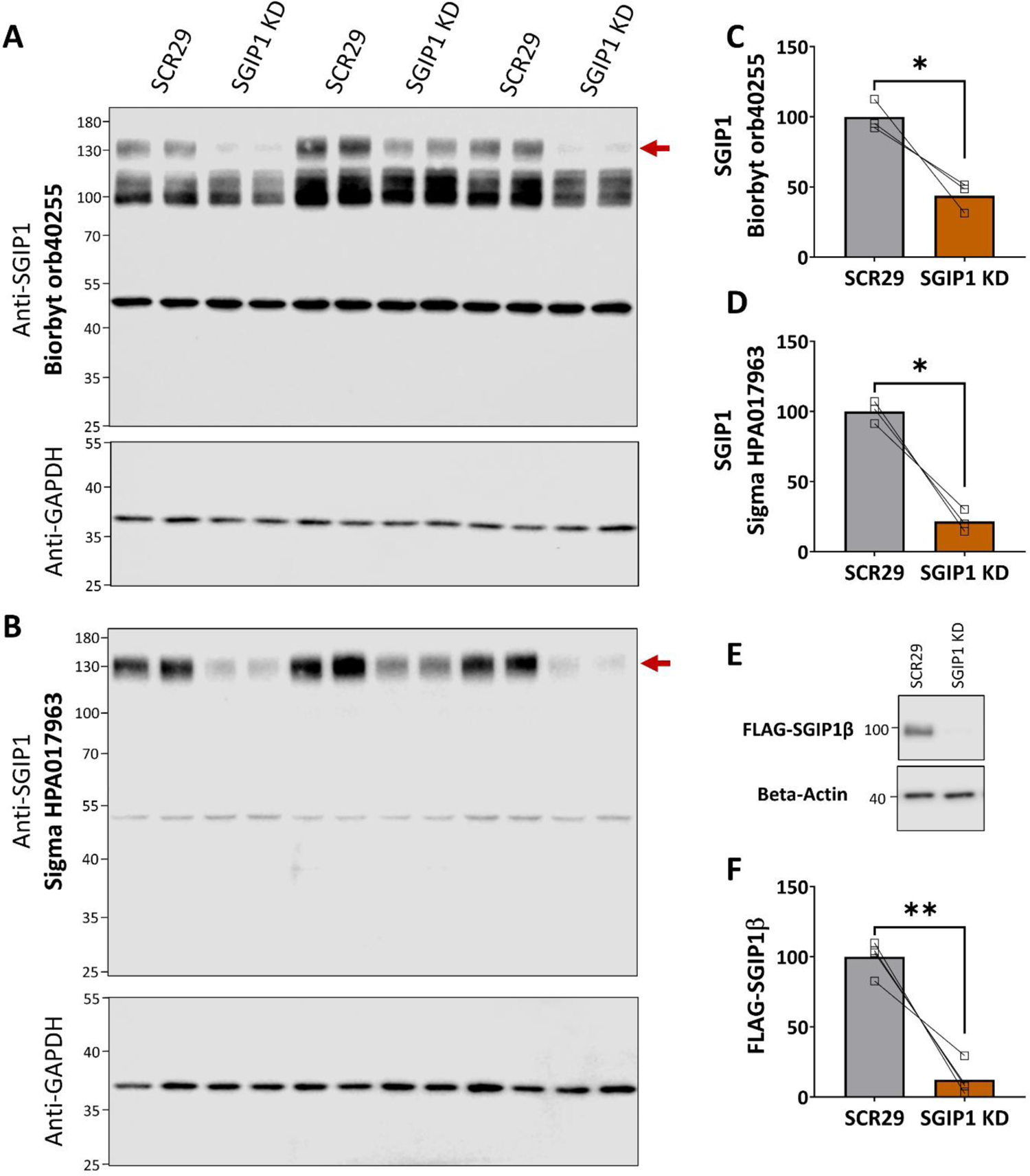
Validation of SGIP1 shRNA and antibodies. (**A-B**) Immunoblots of DIV14 cortical neuronal samples lentivirally transduced with SGIP1 shRNA or a 29mer scrambled control. Two different anti-SGIP1 antibodies were tested: (**A**) Biorbyt orb40255 and (**B**) Sigma HPA017963. Both antibodies recognised a band at around 130kDa (red arrow) that was reduced in intensity in SGIP1 shRNA samples compared to the 29mer scrambled control (**C**-**D**), suggesting this band is SGIP1 and/or SGIP1α. Biorbyt orb40255 also recognises a non-specific smear from around 100kDa to 120kDa and a band around 50kDa. Sigma HPA017963 also faintly recognised a band around 50kDa. This could represent a non-specific band or possibly a short isoform of SGIP1 or closely related protein that is insensitive to our shRNA. SGIP1b was cloned out of our cortical neurons, suggesting that it may be one of the principal isoforms found in these cells, and FLAG-SGIP1b runs around 100kDa (see **E**). Sigma HPA017963 does not recognise a band at that molecular weight. However, this is likely because the antigen used to generate this antibody (residues K254-D345) almost completely overlaps with the deletion found in SGIP1b (N273-E438). The epitope used to generate Biorbyt orb40255 (M1-R30) is present in both SGIP1 and SGIP1b, but the non-specific smear from 100kDa to 120kDa likely obscures this band. For clarity, all experiments in Fig. 3 used the Sigma HPA017983 as it produced fewer non-specific bands, with the assumption that all isoforms of SGIP1 would be knocked down in a similar ratio. Blots were stripped and reprobed for GAPDH as a loading control. (**C-D**) Paired t-tests. n = 3 independent experiments. (E) Representative immunoblots showing shRNA knockdown (KD) of overexpressed FLAG-SGIP1b and a beta-actin housekeeping control in HEK293T cells. FLAG-SGIP1b and a 25-mer SGIP1-targeting shRNA or a 29-mer non-targeting control (SCR29) shRNA were co-transfected into HEK293T cells and left for 3 days. (F) Quantification of data represented in (**E**). The 25mer shRNA targeting SGIP1 knocked down overexpressed FLAG-SGIP1b by about 88% compared to a scrambled (non-targeting) 29mer control (SCR29). Paired t-test. n = 4 independent experiments.

**Supplementary Figure 2:**
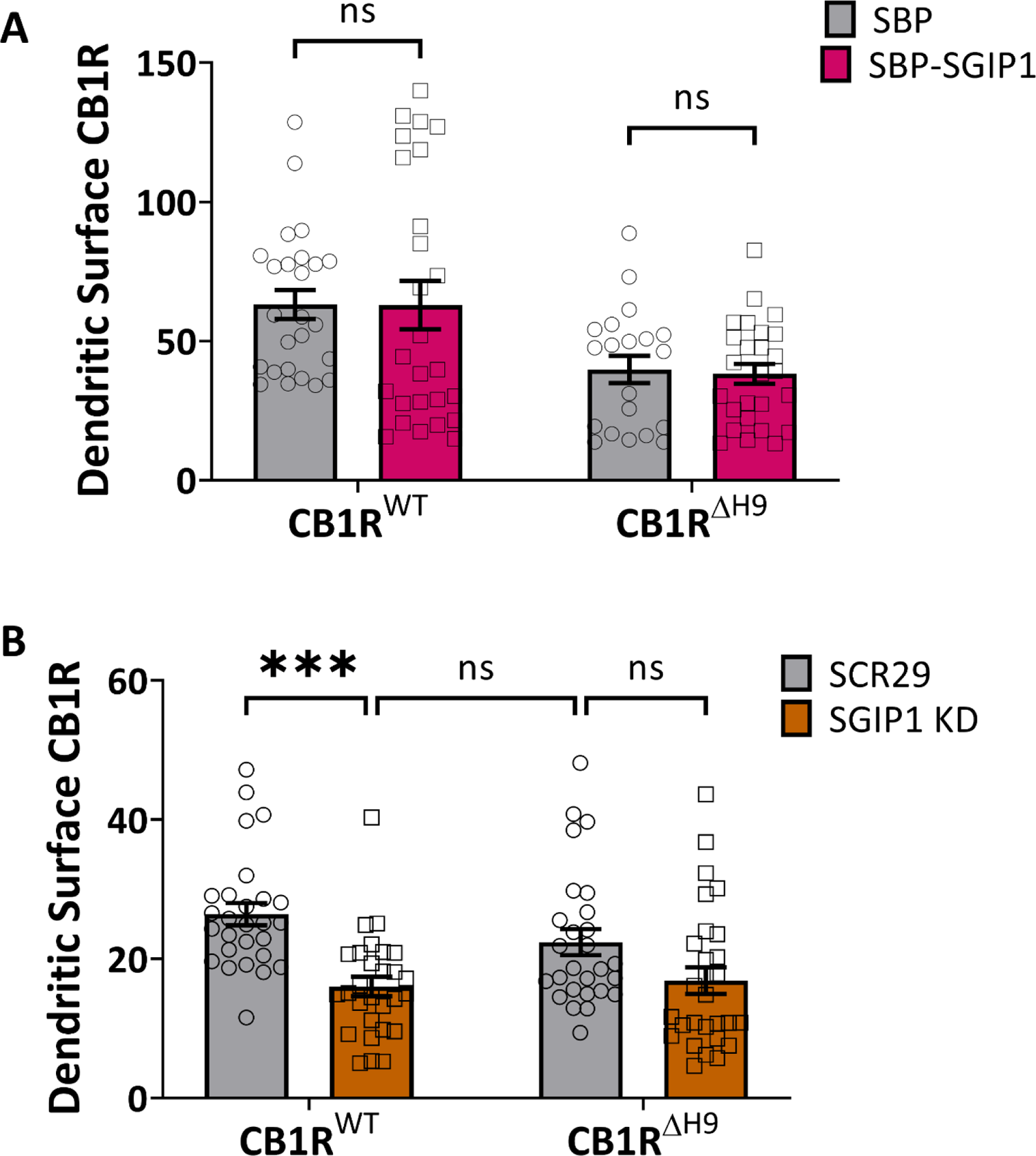
SGIP1 and dendrites. (A) Quantification of data represented in Fig. 2A. Overexpression of SGIP1b isoform has no effect on dendritic surface levels of either CB1R^WT^ or CB1R^ΔH9^. Surface fluorescence was normalised to total fluorescence and shown as a percentage of axonal CB1R^WT^/SBP control (see Fig. 3B). Two-way ANOVA with Sidak’s *post hoc* test; n = 19-27 neurons per condition from four independent neuronal cultures. (B) Quantification of data represented in Fig. 3A. Knockdown of SGIP1 reduces dendritic surface levels of CB1R^WT^, but not CB1R^ΔH9^, suggesting that an isoform other than SGIP1b may affect CB1R surface expression in dendrites. Surface fluorescence was normalised to total fluorescence and shown as a percentage of axonal CB1R^WT^/SCR29 (see Fig. 3B). Two-way ANOVA with Sidak’s *post hoc* test. N = 27-28 neurons from five independent neuronal cultures per condition.

**Supplementary Figure 3:**
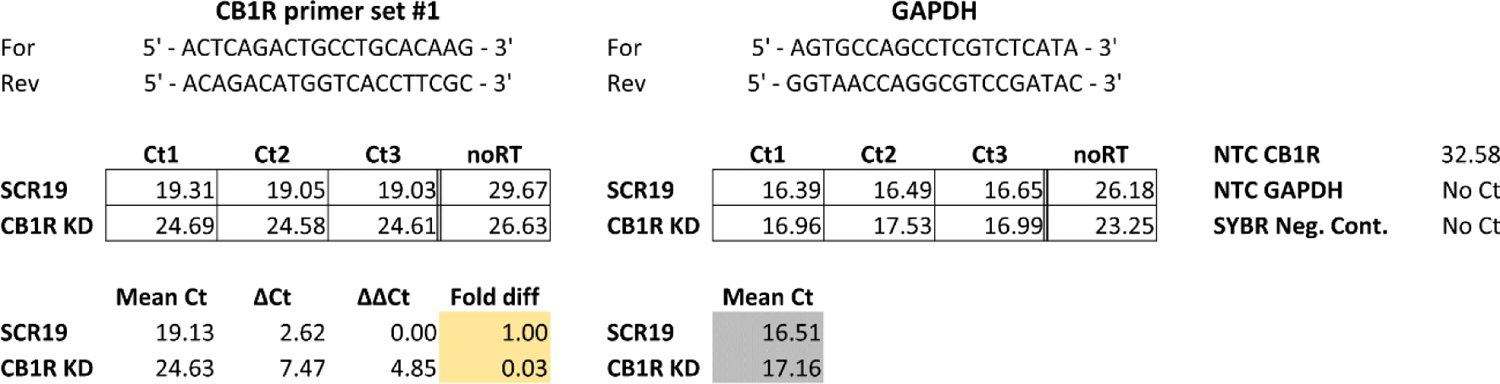
qPCR primers were validated against KD samples. Specificity of CB1R qPCR primer set used in Fig. 4G was tested against DIV14 cortical neuronal samples lentivirally transduced with CB1R shRNA (target sequence from (43) or a 19mer scrambled control (target sequence from (44)). There was a 97% reduction in transcript levels in CB1R KD neurons compared to SCR19 control, suggesting that the primers used are specific for CB1R mRNA.

